# Filtration extraction method using microfluidic channel for measuring environmental DNA

**DOI:** 10.1101/2021.11.18.468931

**Authors:** Takashi Fukuzawa, Yuichi Kameda, Hisao Nagata, Naofumi Nishizawa, Hideyuki Doi

**Affiliations:** GO!FOTON INC., Tsukuba, Japan; Biryu Planning, Tokyo, Japan; Department of Anthropology, National Museum of Nature and Science, Tsukuba, Japan; Graduate School of Information Science, University of Hyogo, Kobe, Japan

**Keywords:** eDNA, extraction, filtering, microfluidic

## Abstract

The environmental DNA (eDNA) method, which is widely applied for biomonitoring, is limited to laboratory analysis and processing. In this study, we developed a filtration/extraction component using a microfluidic channel, Biryu-Chip (BC), and a filtration/extraction method, BC method, to minimize the volume of the sample necessary for DNA extraction and subsequent PCR amplification. We tested the performance of the BC method and compared it with the Sterivex filtration/extraction method using aquarium and river water samples. We observed that using the BC method, the same concentration of the extracted DNA was obtained with 1/20–1/40 of the filtration volume of the Sterivex method, suggesting that the BC method can be widely used for eDNA measurement. In addition, we could perform on-site measurements of eDNA within 30 min using a mobile PCR device. Using the BC method, filtration and extraction could be performed easily and quickly. The PCR results obtained by the BC method were similar to those obtained by the Sterivex method. The BC method required fewer steps and therefore, the risk of DNA contamination could be reduced. When combined with a mobile PCR, the BC method can be applied to easily detect eDNA within 30 min from a few 10 mL of the water sample, even on-site.

## Introduction

Environmental DNA (eDNA) in aquatic environments has been used to detect the distribution of the species (Doi et al., 2017a; Takahara, Minamoto, Yamanaka, Doi, and Kawabata, 2012; Katano, Harada, Doi, Souma, and Minamoto, 2017). eDNA has been detected in water from various ecosystems, including streams, lakes, ponds, reservoirs, canals, lakes, and oceans (Doi et al., 2017; Takahara, Minamoto, Yamanaka, Doi, and Kawabata, 2012; Katano, Harada, Doi, Souma, and Minamoto, 2017; Fornillos et al., 2019). eDNA measurements have been mainly performed using quantitative real-time PCR (qPCR) (Doi et al., 2017a; Takahara, et al., 2012; Katano et al. 2017; Fornillos et al., 2019). However, it is limited to laboratory analysis, which usually takes several hours. These time delays often limit the range of uses for on-site eDNA detection (Thomas et al., 2020; Nguyen et al., 2018). Field-portable DNA extraction and PCR platforms offer a way for species detection by eDNA analysis on-site (Thomas et al., 2020; Nguyen et al., 2018; Thomas, Howard, Nguyen, Seimon, and Goldberg, 2018). However, this approach takes a similar time as that of the laboratory measurements. Recently, Doi et al. (2021) reported a simpler extraction method in the field using PicoGene PCR1100 (mobile qPCR; Nippon Sheet Glass, Sagamihara, Japan), which allowed eDNA detection in a relatively short time (Doi et al., 2021). This method required approximately 30 min to complete after sample water collection for DNA measurement by qPCR. However, time and facilities are limited for sample analysis in the field, and there is a need for a simpler and more efficient pretreatment method, including filtration. In particular, the simplicity of species-specific eDNA measurement should be one of the key features to allow for the large-scale implementation of eDNA technology.

In eDNA measurements, the sensitivity of DNA detection is important. Therefore, it is essential not to compromise on the concentration and yield of the extracted DNA from the water samples. Although various extraction methods using DNA isolation kits such as the DNeasy Blood and Tissue Kit (Qiagen, Hilden, Germany) (Tsuji et al., 2019), PowerWater (Hinlo, Gleeson, Lintermans, and Furlan, 2017; Coster, Dillon, Moore, and Merovich, 2021), and PowerSoil (Sakata et al., 2020; Eichmiller, Miller, and Sorensen, 2016; Díaz et al., 2020) have been used, it is necessary to simplify the process from water sampling to eDNA extraction for field measurements and general users.

They include disc-shaped filters (e.g., GF/F glass filters) (Tsuji et al., 2019), nucleopore filters (e.g., 0.2-μm pored filter) (Takahara et al. 2012; Sassoubre, Yamahara, Gardner, Block, and Boehm, 2016), and cartridge filters (e.g., Sterivex) (Miya et al., 2016). However, if less volume of the DNA extraction solvent is used, it is difficult to contact the entire area of the filter uniformly. As a result, DNA would partially remain on the filter. Consequently, the DNA yield would decrease. In contrast, when a microfluidic channel is used for filtration, the entire surface of the filter on the channel comes in contact with the DNA extraction solvent. Accordingly, DNA can be extracted efficiently and a high concentration of the extracted DNA can be obtained.

In this study, we developed a filtration/extraction component using a microfluidic channel to minimize the volume of the sample necessary for DNA extraction, simultaneously yielding a high concentration of eDNA for subsequent PCR amplification. We demonstrated that eDNA can be measured easily with low contamination risk using this component (Muha, Robinson, Garcia de Leaniz, and Consuegra, 2019). We also tested the performance of the developed microfluidic channel using samples from aquarium and river water and compared the results with conventional methods using Sterivex filters, DNeasy Blood and Tissue Kit, and benchtop qPCR for various fish species.

## Materials and Methods

### Material design for microfluidic channel chip: Biryu-Chip

We developed a filtration/extraction chip with a microfluidic channel, called the Biryu-Chip (Biryu: micro-channel in Japanese) and the filtration/extraction method is termed as the Biryu-Chip (BC) method. A schematic diagram of the BC is shown in Fig. 1a, and a photograph is shown in Fig. 1b. The base material was made of cycloolefin polymer and molded by injection molding. The filter was made of polyvinylidene difluoride, the same material as the Sterivex filter. The upper channel of the filter is 1 mm wide (i.e., the effective width of the filter), 0.5 mm high, and 40 mm long, with a cross-sectional area of 0.5 mm^2^. During filtration, the sample was passed from the inlet to the outlet using a syringe and the eDNA was trapped on the filter (Fig. 1c). Finally, the DNA was extracted into the buffer (Fig.1d).

**Fig 1.**
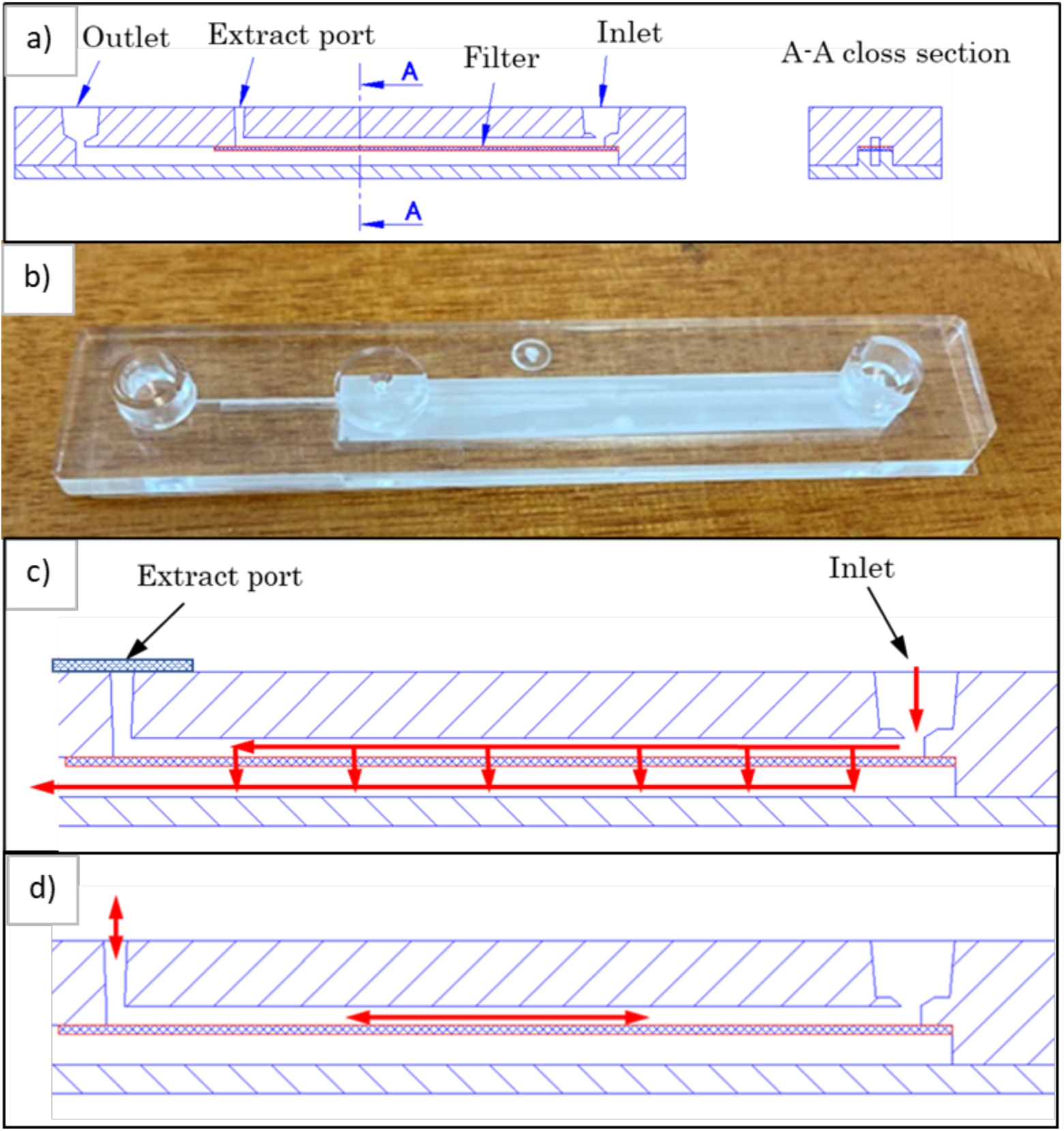
a) Schematic diagram of the Biryu-Chip and b) Biryu-Chip, c) Sample flow at filtering, and d) The flow of extraction solvent during the extraction.

### Validation experiments

Four experiments were performed to validate the BC method. Experiment 1: determining the volume of DNA extraction solvent for the BC method by a benchtop qPCR using water samples spiked with rainbow trout (*Oncorhynchus mykiss*) aquarium water; 2: comparing the BC method with the Sterivex method by a benchtop qPCR analysis (hereafter referred to as Sterivex-qPCR) using water samples spiked with rainbow trout aquarium water; 3: field validation of the BC method performed using the river water samples and compared with the results using the Sterivex method by detecting sweetfish (*Plecoglossus altivelis*) DNA on a benchtop qPCR; and 4: field validation of the BC method using a mobile PCR (hereafter referred to as BC-mobile PCR) was done from another sample of river water which was a habitat to the common carp (*Cyprinus carpio*). The outline of the extraction steps of the Sterivex and BC methods is shown in Fig. 2.

**Fig. 2.**
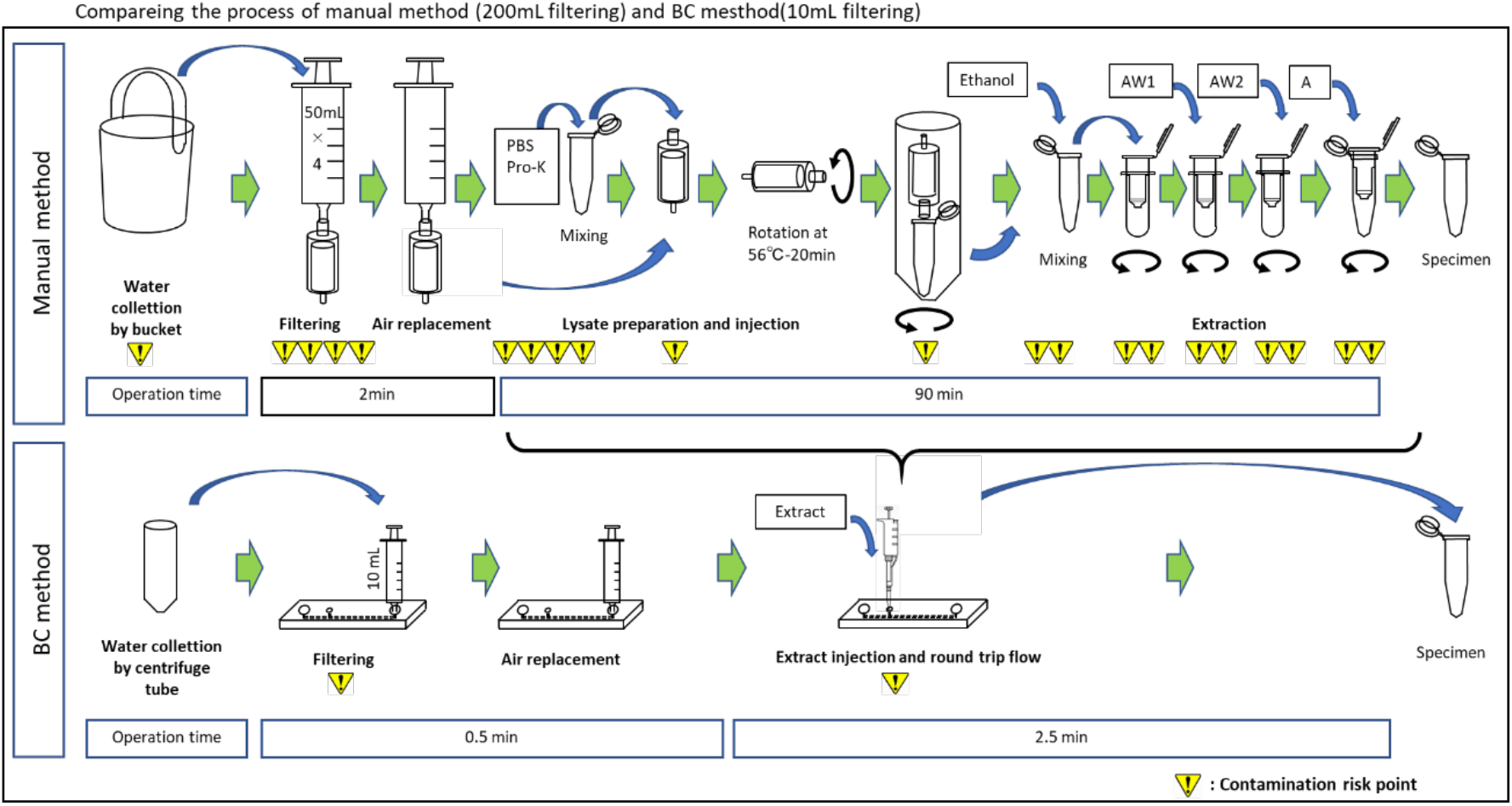
Comparison of work flow between the method of Miya et al. (2016) and the BC method.

### Experiment 1

We collected 4 mL of rainbow trout aquarium water and made up the final volume to 200 mL with pure water, mixed well, and then filtered 10 mL of diluted sample each through eight BCs. Extraction of the DNA was performed according to the BC method. Different volumes of the extraction solvent, as explained later, were used for DNA extraction in increasing volume, that is 6, 8, 10, 12, 14, 16, 18, and 20 μL representing eight levels. We measured the concentration of the extracted DNA using a benchtop qPCR, as described below.

### Experiment 2

We collected 2 mL of rainbow trout aquarium water and made up the volume to 250 mL with pure water and mixed well. Then, 10 mL of the diluted sample was filtered by the BC filter and 200 mL by the Sterivex filter. DNA extraction from the BC filter was done using 20 μL of the extraction solvent. DNA from the Sterivex filter was extracted using a DNeasy Blood and Tissue Kit. We measured the concentration of DNA extracted by both methods using a benchtop qPCR.

### Experiment 3

We collected 250 mL of surface river water from the middle basin of the Sagami River (35.575099°N, 139.308802 °E), a known sweetfish habitat. Two water samples (10 mL each) were filtered by two BC filters and 200 mL was filtered by the Sterivex filter. DNA from the BC filter was then extracted with 10 and 20 μL of the extraction solvent. DNA from the Sterivex filter was extracted using a DNeasy Blood and Tissue Kit. We measured the concentration of DNA extracted by the BC and Sterivex methods using a benchtop qPCR.

### Experiment 4

In this experiment, we used eDNA of the common carp from the Sakai River. We collected water samples at several points (from bridge 1: 35.600385°N, 139.354699°E to bridge 6: 35.604273°N, 139.335399°E) in the Sakai River (Fig. 3) using a centrifuge tube attached to a fishing line. The sample water (10 mL) was filtered through the BC chip, resulting in an extraction volume of 20 μL.

**Fig. 3.**
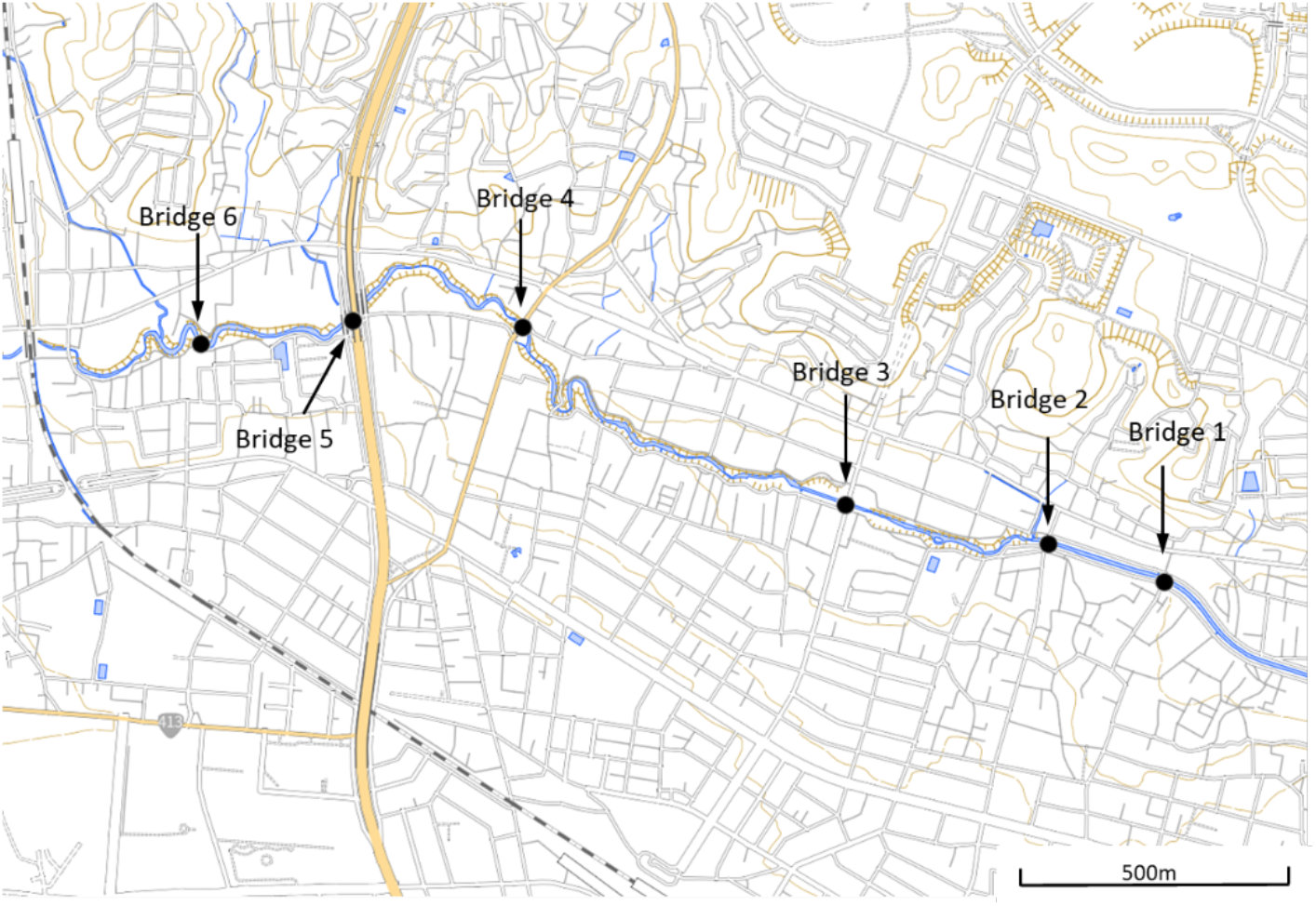
Sampling points in the Sakai River for Experiment 4 created by processing the GSI tiles (Map data from Geospatial Information Authority of Japan (GSI) with permission: https://maps.gsi.go.jp/development/ichiran.html).

### Filtration and DNA extraction by the BC method

We performed sample filtration and DNA extraction by the BC method using the setup shown in Fig. 4. The procedure followed is given below:

1. Water was collected using a disposable container, such as a 50 mL centrifuge tube. A syringe (e.g., 10 mL volume) was used to aspirate a predetermined volume of water and then the water sample was injected into the BC inlet and filtered.
2. The syringe was removed after sample filtration. The plunger was pulled to draw air and then the syringe was inserted into the BC inlet. Then, the plunger was pushed to drain the excess water from the chip.
3. The seal attached to the extraction port of the BC filter was removed. The extraction solvent was then injected into the filter (10 and/or 20 μL for Experiment 2–4, 6–20 μL for Experiment 1) and mixed with a pipette to allow the movement of the extraction solvent throughout the upper channel of the filter for 2 min. Solution A of the Kaneka Easy DNA Extraction Kit (Version 2) (Kaneka, Tokyo, Japan) was used as the extraction solvent.
4. The extraction solvent was drawn from the extraction port using a pipette and then transferred to a microcentrifuge tube to obtain the extracted DNA sample.

**Fig. 4.**
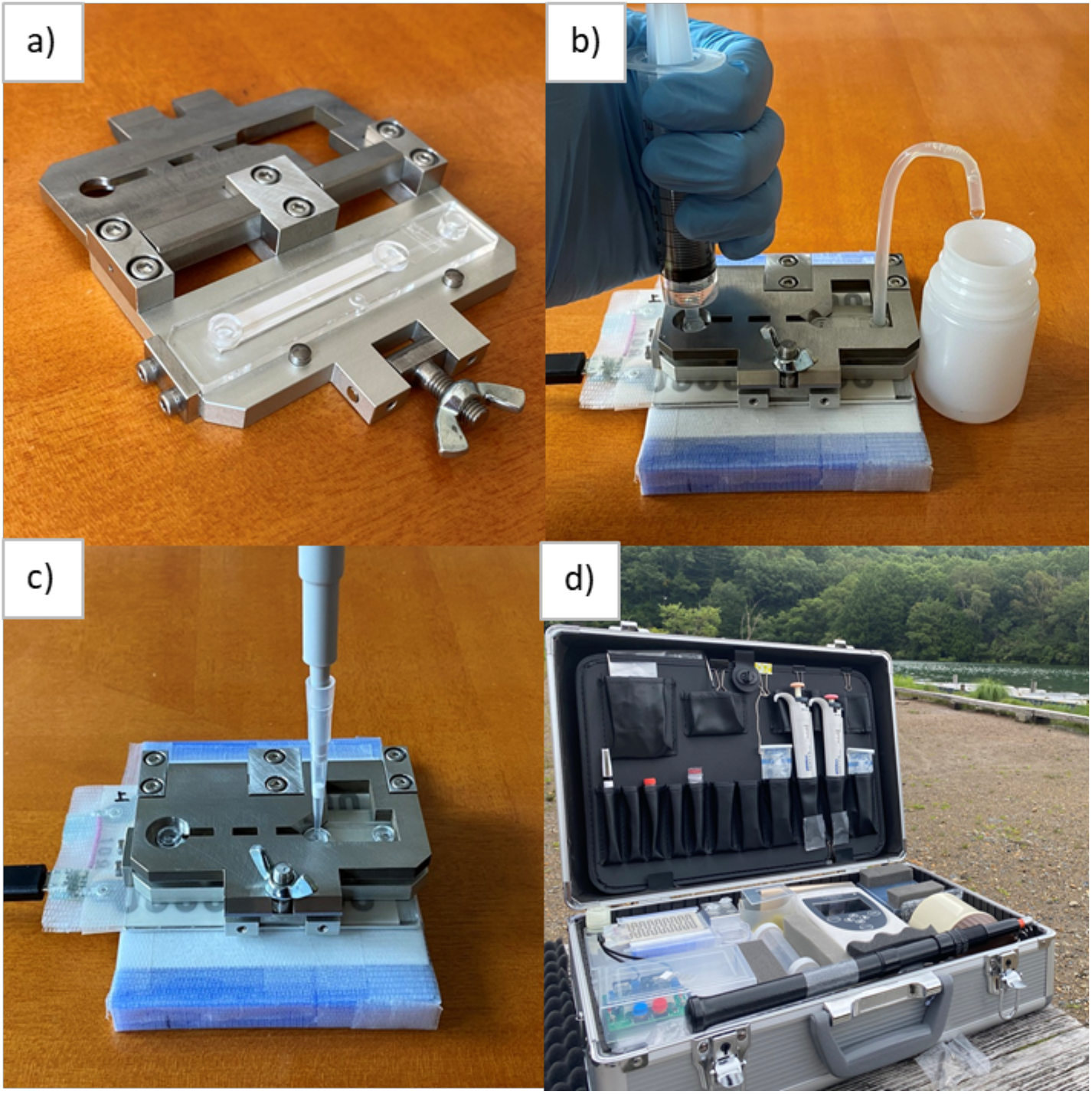
a) Tool for filtering and extraction using the Biryu-Chip, b) Filtering process, c) Extraction process, and d) eDNA measurement Kit (440 × 310 × 128 cm case).

### DNA extraction by the conventional method

To evaluate the performance of the BC method in Experiments 2 and 3, we compared its results with that of the conventional method, which is a combination of the filtration method using the Sterivex filter (Takahara, Minamoto, Yamanaka, Doi, and Kawabata, 2012; Sassoubre, Yamahara, Gardner, Block, and Boehm, 2016) and the extraction method using the DNeasy Blood and Tissue Kit (Tsuji et al., 2019). Filtration using the Sterivex filter is relatively easy to perform on-site without the requirement of sophisticated infrastructure and the DNA extraction method using the DNeasy Blood and Tissue Kit is a widely used method in academic research (Minamoto et al., 2021). Filtration and DNA extraction from the water samples collected in the experiments above were performed according to the manufacturer’s instructions (Minamoto et al., 2021).

### PCR kit used for benchtop qPCR and mobile PCR

It is known that the efficiency of PCR amplification decreases in the presence of substances such as humic acid, which is a PCR inhibitor (Uchii et al., 2019). Since the purification of the water sample was not involved in the BC method, the effect of PCR inhibitors may be significant. Therefore, in this study, we used KAPA3G Plant PCR Kit (Sigma-Aldrich, Missouri, USA) which is relatively resistant to PCR inhibitors.

### Benchtop qPCR

Quantification of eDNA from all the samples except those in Experiment 4 was done using the StepOnePlus Real-Time PCR System (Thermo Fisher Scientific, Massachusetts, USA). For the laboratory qPCR analyses, we used the same set of primer-probe as in the on-site measurements. Similarly, the PCR template mix used was as described in the previous studies (Doi et al., 2017a, Uchii et al., 2019). Each TaqMan reaction contained 900 nM of each primer (forward and reverse), 125 nM TaqMan-Probe, 0.03 U/μL qPCR master mix (KAPA3G Plant PCR Kit), and 1.5 μL of the eDNA solution. The final volume for a single PCR assay was made up to 15 μL by adding distilled water (DW). The volume of the eDNA solution added was 10% of the volume of the PCR mixture. The qPCR conditions were as follows: 95 °C for 20 s, followed by 55 cycles of 95 °C for 4 s and 60 °C for 20 s. Three replicates were performed for each sample and no-template control (NTC). Standard curves of qPCR measurements had R^2^ = 0.995–0.998 and PCR efficiency = 95.4%–105.2%.

### Mobile PCR

Before sampling, we prepared a PCR pre-mix with preliminary mixing of the master mix and primer probe. This pre-mix was brought on-site for PCR analysis. Each TaqMan reaction contained 900 nM of each primer (forward and reverse), 400 nM of TaqMan-Probe, 0.1 U/μL qPCR master mix (KAPA3G Plant PCR Kit). The final volume for PCR pre-mix was made up to 14.4 μL by adding DW. Then, 1.6 μL of the eDNA solution was added to the tube containing the pre-mix. The volume of the eDNA solution added was 10% of the volume of the PCR mixture. This was followed by the injection of this solution into the flow path of the PicoGene PCR1100. The PCR pre-mix was stored in a cooler at 5 °C until PCR was performed. The mobile PCR conditions were as follows: 95 °C for 15 s, followed by 50 cycles of 95 °C for 3.5 s, and 60 °C for 10 s. Additionally, to check the cross-contamination in the PCR machine on-site, we performed an NTC using DW after completion of all the PCR measurements for the day (PCR control) on site. The primer-probe sets are listed in Table 1.

**Table 1.**
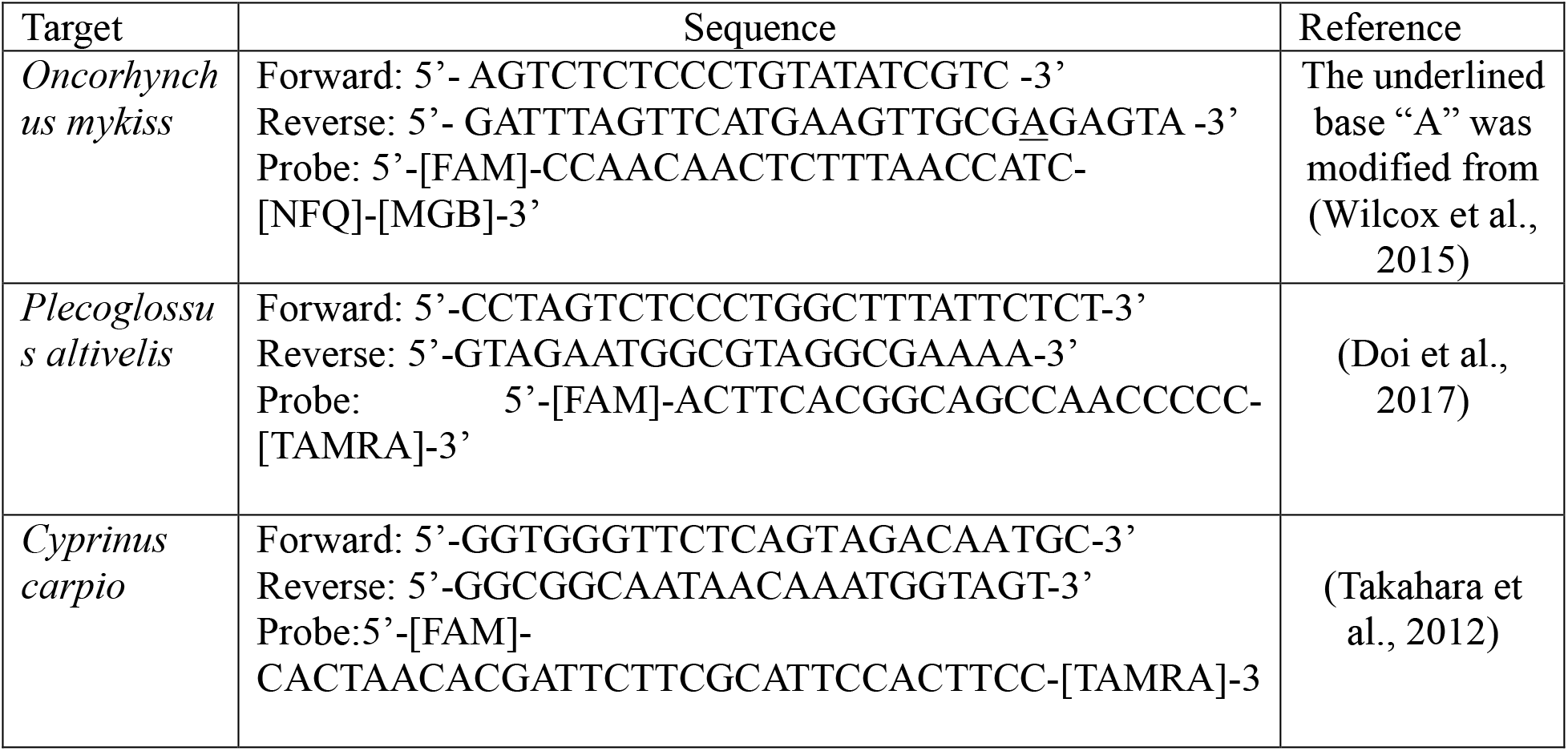
Primer-probe sets using this study.

### Statistical analysis

All statistical analyses were performed using the R package (ver. 4.1.1) (R Core Team, 2021) and the ggplot2 package. The significance level was set to α = 0.05. We performed a simple linear model to predict the DNA extraction volume of the BC method and the DNA concentration using the “lm” function. For categorical data of Experiments 2 and 3, we performed one-way ANOVA using the “aov” function. When the ANOVA was significant, we performed the post-hoc test by Tukey’s multiple comparisons using the “TukeyHSD” function for comparing each difference.

## Results

### Experiment 1: Determination of the volume of the extraction solvent

Fig. 5 shows DNA extraction by the BC method and qPCR results (DNA copy number) using different volumes of the extraction solvent. From the linear model results, we observed a significant relationship between the volume of the extraction solvent and DNA copy number (R^2^ = 0.9167, P for R^2^ and slope < 0.0001).

**Fig. 5.**
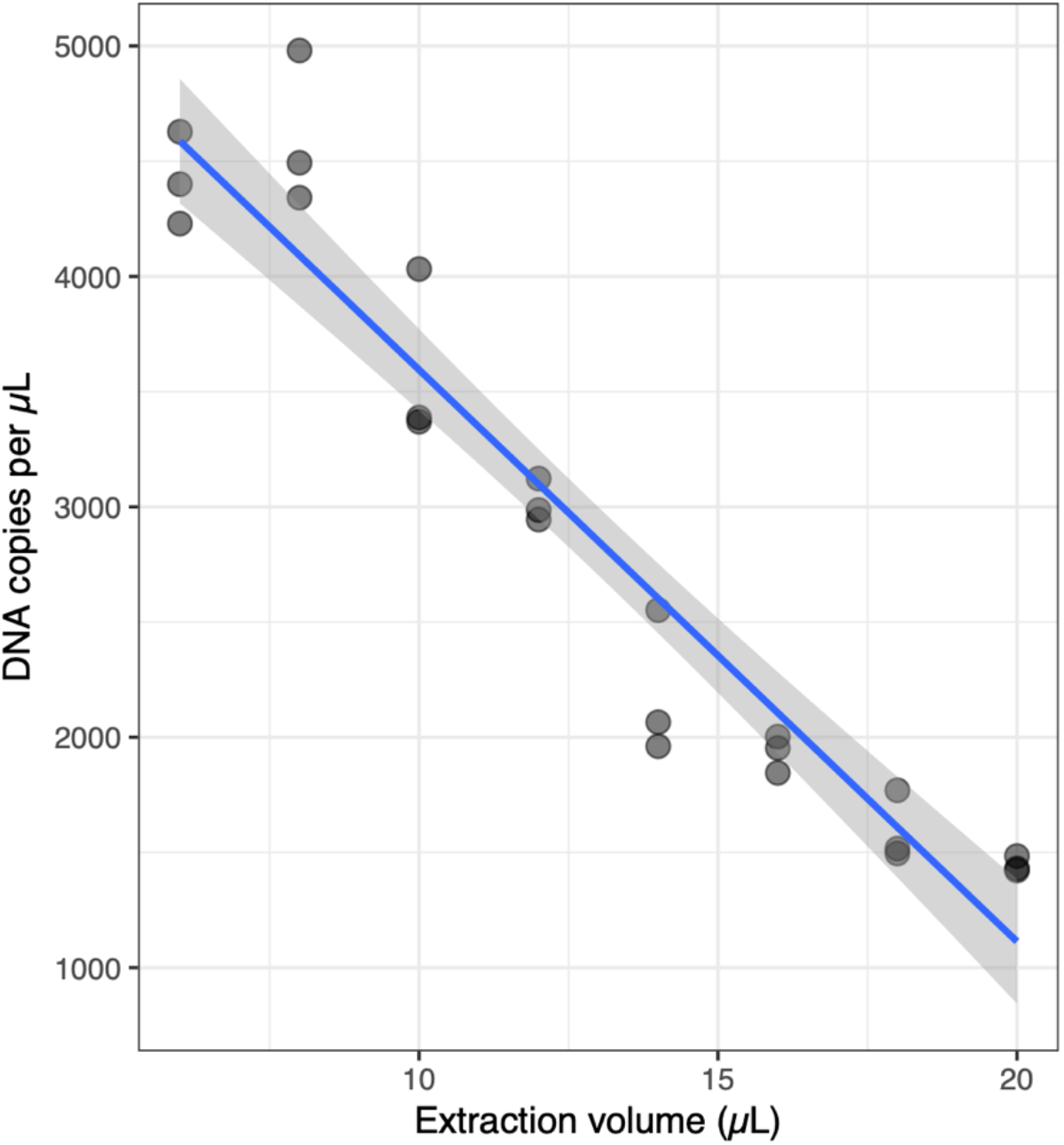
The relationship between DNA extraction volume and measured DNA copy number of the rainbow trout in Experiment 1. The line and grey shadow represent linear regression and the 95% CI of the slope, respectively. The dot represents each data point.

### Experiment 2: Comparison using the samples spiked with aquarium water

Fig. 6 shows the qPCR results of the amplification of DNA from the rainbow trout in the samples extracted by the Sterivex and BC methods. We observed significant differences in ANOVA (F= 11.65, P < 0.0001) and Tukey’s comparison (P < 0.05) for the pair of Sterivex 1 and BC 1 (Fig. 6). The average copy number of the four extracted samples were 385 ± 141 copies μL^-1^ for the Sterivex method and 599 ±154 copies μL^-1^ for the BC method. The coefficient of variation (CV: standard deviation/mean value) of the Sterivex and BC method were 0.33 and 0.26, respectively, indicating that the BC method did not show a large variability. These results suggest that the eDNA concentration extracted using the BC method from various samples was similar despite a lower filtration volume (5%). In the negative control, no rainbow trout DNA was detected in the extracted sample of non-spiked water in each method.

**Fig. 6.**
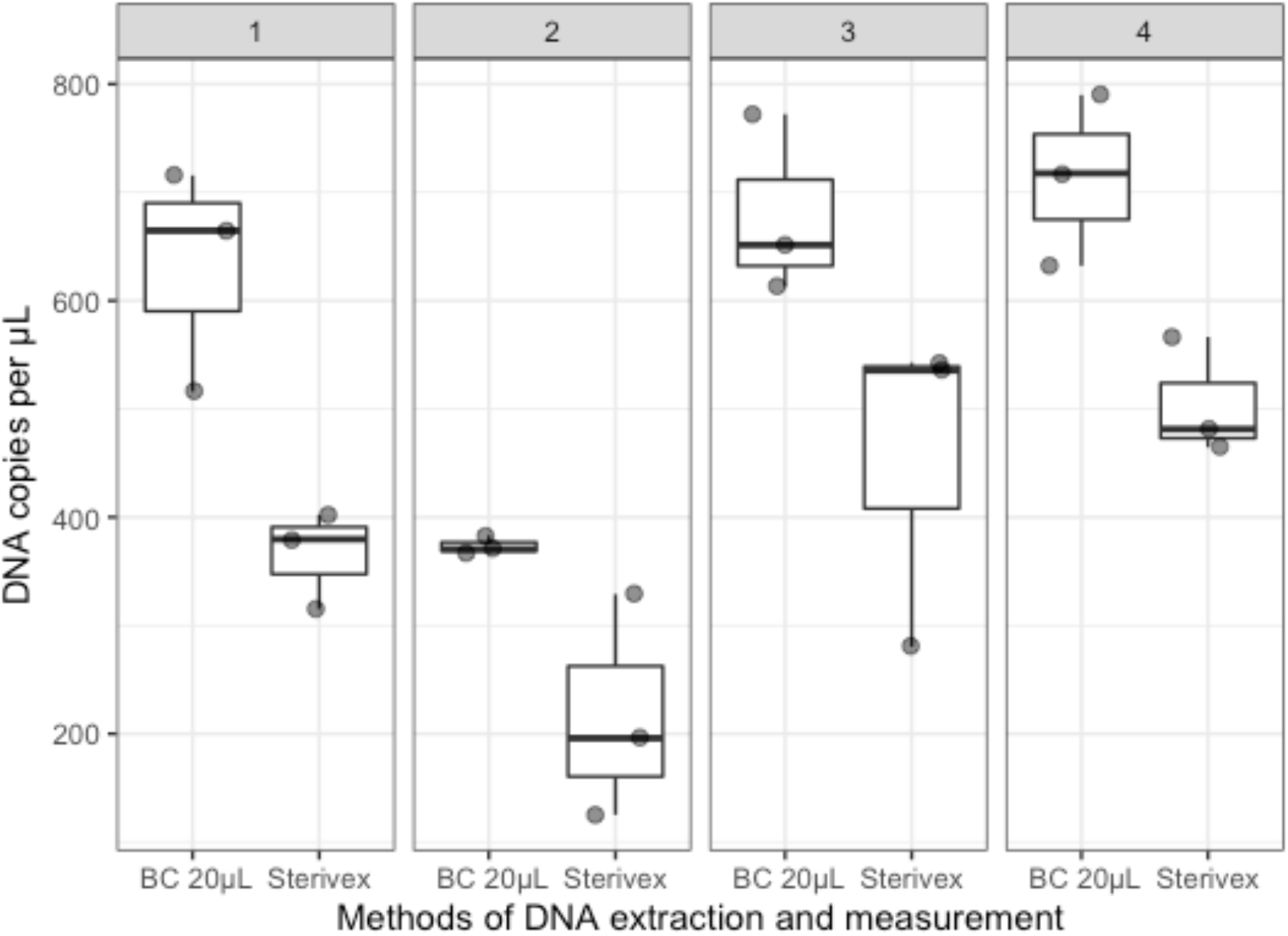
Boxplot for Experiment 2. The boxes indicate ± 25% quartiles with the median (bar) and the bars indicate ± 1.5x quartiles. The dot represents each data point.

### Experiment 3: Comparison with filed samples

The water of the Sagami River was used in this experiment. We observed differences in the concentration of DNA from the sweetfish as determined by the Sterivex and the BC method using 10 and 20 μL of extraction solvents (Fig. 7). We also observed that ANOVA for the three categories was significant (F = 55.66, P < 0.0001). The BC method using 10 μL of the extraction solvent had significantly highest DNA concentration as determined by the Tukey’s test (P < 0.0001). The mean value of the copy number from extracted samples for each category was 34 ±6.7 copies μL^-1^ for the Sterivex method, 38 ± 12.3 copies μL^-1^ for the BC method with 20 μL of the extraction solvent, and 82 ± 16.5 copies μL^-1^ for the BC method with 10 μL of the extraction solvent. The CV values of the Sterivex method, BC method with 20 μL and 10 μL of the extraction solvent were 0.18, 0.30, and 0.19, respectively.

**Fig. 7.**
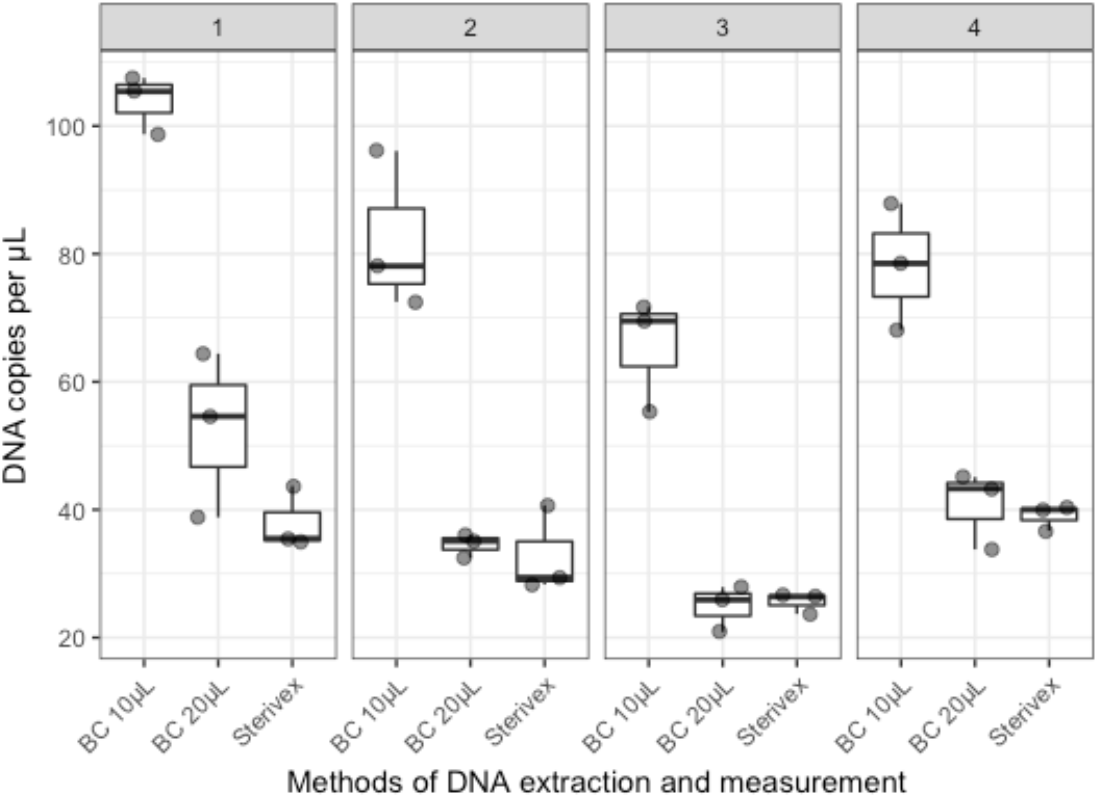
Boxplot for Experiment 3. The boxes indicate ± 25% quartiles with the median (bar) and the bars indicate ± 1.5x quartiles. The dot represents each data point.

### Experiment 4: eDNA measurement in the field

Fig. 8 shows the Ct values of carp eDNA obtained by the on-site BC-mobile PCR methods. We detected high concentrations of the common carp DNA at all survey sites. From these results, it could be inferred that the common carp was living in river water between the bridges 5 and 6.

**Fig. 8.**
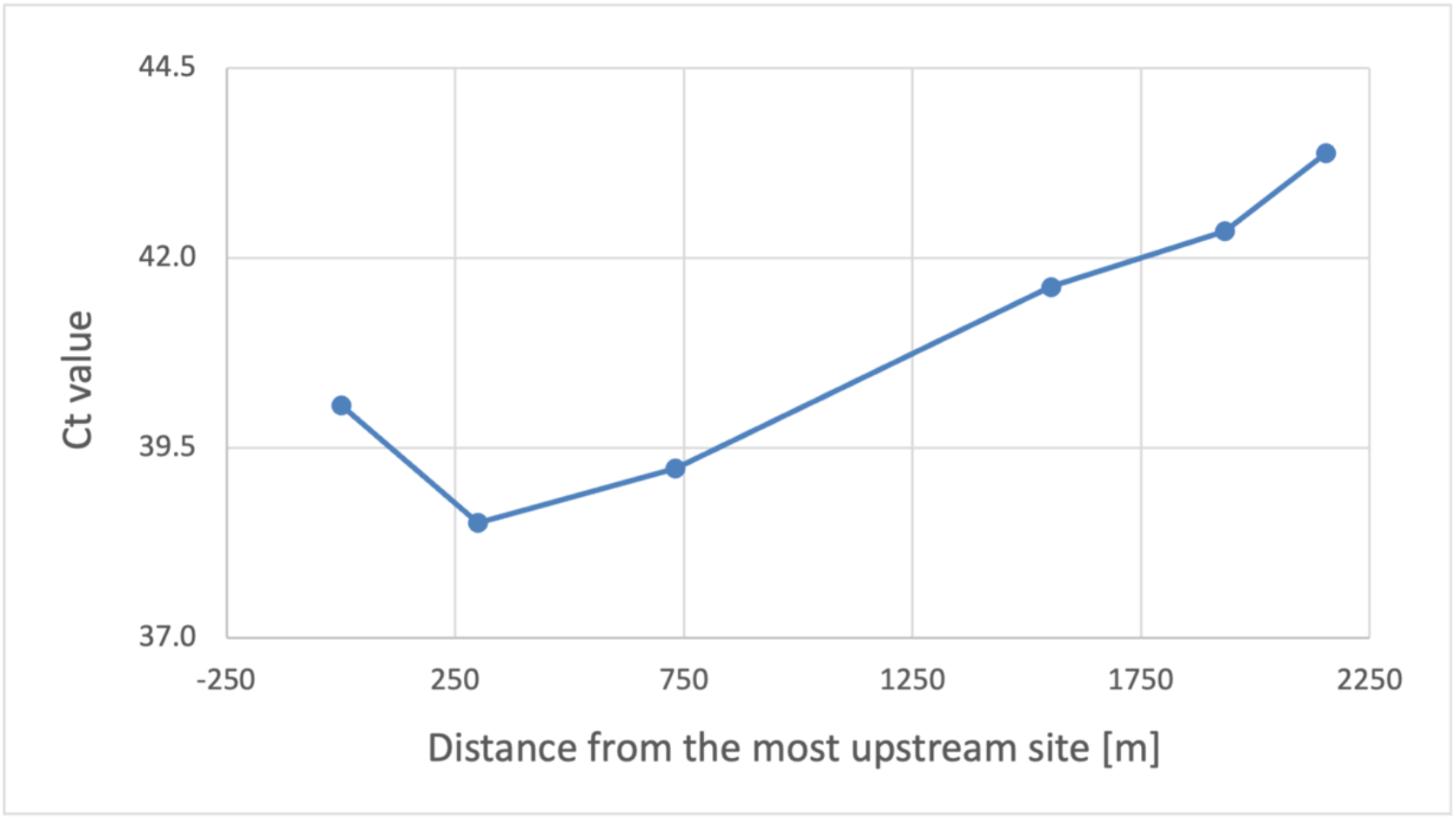
The relationship between distance from the most upstream site and Ct values of the carp DNA.

## Discussion

In this study, we developed a new filtration/extraction method called the BC method for eDNA measurements from various water ecosystems. We observed that using this method, the concentration of DNA from the extracted samples was almost similar to the concentration of DNA extracted using the conventional methods even with 1/20–1/40 of the filtration volume that is generally used in the Sterivex method. Therefore, the BC method can be used to obtain a highly concentrated DNA sample even with low filtration volumes. Although the efficiency of the BC method may change according to the target eDNA, it is not inferior to the conventional methods in terms of the DNA yield, suggesting that it can be widely used for eDNA measurement.

As the filtering volume of BC method is small while measuring eDNA at lower concentrations, the volume of eDNA to be filtered is also small. As a result, the variation may be larger than when using a large volume of sample water which may affect the detection limit. This problem with the detection limit could be solved by further increasing the concentration ratio by optimizing the filtration and extraction conditions of the chip. In addition, even if the detection limit is reduced, it can be expected that the BC method has industrial applications where it can be used by balancing the merits of simplification.

In the BC method, the volume required for filtration can be can be greatly reduced, and the method makes water sampling simple. For example, in the design of the BC method, it requires less than 25 mL filtration volume compared to 0.5 to a few liters of water for the previous methods (Tsuji et al., 2019). For measurements of eDNA from the habitats where it is difficult to collect water, such as small wetlands (Doi et al., 2017b), the use of the BC method would be preferable. The process after filtration in the BC method is much simpler than the Sterivex method and does not require a high level of skill. The working time was reduced from 90 min to 3 min. In addition, special equipment such as centrifuges and rotators are not required, which makes it easier to introduce measurement equipment in the process. As a result of the simplification of filtration and extraction, a set of components and instruments from water sampling to PCR can be stored in an attached case (Fig. 4d), and eDNA detection can be easily performed on-site in approximately 30 min, like our previous method (Doi et al. 2021).

The situations in which these advantages are of high importance include on-site eDNA measurement and in study areas where it is difficult to collect water such as wetlands and valleys. The BC-mobile PCR method has the following benefits: measurements at multiple points in a day and the sampling site which is difficult to access.

The filtration and extraction steps of the eDNA measurement has the risk of contamination. Moreover, the risk is also higher if there are several reusable components, such as filtering and extraction equipment, including a filter funnel and centrifuge machine. However, the BC method uses disposable materials, including the chip, thus reducing the contamination risk in the eDNA measurements. In addition, since filtration and extraction can be performed consistently in the closed space of the BC, there are fewer opportunities for the samples and reagents to be exposed to the surrounding air. This further reduces the risk of contamination compared to other filtration and extraction methods. Furthermore, the principal of the BC method can be used for detecting bacteria or viruses by changing the design of the BC and filter. Thus, the BC method can be applied to broad scientific fields of research, such as the human health and food sciences.

In conclusion, we developed the Biryu-Chip, a filtration/extraction component, and a method for measuring eDNA using a microfluidic channel. Using the BC method, filtration and extraction can be performed easily and quickly, and it was demonstrated that similar results can be obtained with approximately 1/20–1/40 of the filtration volume compared to the Sterivex method. Since the BC method requires fewer steps, the risk of contamination, which is a problem in eDNA measurements, would be reduced. When combined with a mobile PCR device, this method could detect eDNA within approximately 30 min even on-site. Since a variety of water samples with different turbidity and species are handled in eDNA measurements, it is necessary to validate the usefulness of this method at multiple sites, including samples with more foreign substances, in the future. However, in some cases, it may be necessary to consider the selection of PCR reagents that are more resistant to PCR inhibitors.

## Data availability

All data (Supplementary Table S1) are available in Zenodo (doi: 10.5281/zenodo.5709221).

## Acknowledgments

This study was supported by the Environment Research and Technology Development Fund (JPMEERF20204004).

## Conflict of interest

The commercial affiliations of authors [TF, NN, HN, and YK] do not alter our adherence to the journal policies on sharing data and materials. TF, NN, HN, and YK were employed by the manufacturer of the equipment described. However, none of the authors will directly benefit from the publication of this paper.

